# Enhancement of mitochondrial function fosters B cell immune memory

**DOI:** 10.1101/2022.06.21.497040

**Authors:** Ashima Shukla, Ashutosh Tiwari, Olivier Saulnier, Liam Hendrikse, Bryan Hall, Robert Rickert, Michael D Taylor, Vipul Shukla, Anindya Bagchi

## Abstract

Differentiation of T and B cells to effector and memory cell fates are associated with extensive metabolic changes which are accompanied by altered mitochondrial dynamics. However, whether alterations in mitochondrial structure and function plays an active role in regulating effector versus memory cell fate decisions during immune responses remains unclear. Our studies here characterize changes in mitochondrial dynamics in activated B cells and show that increased mitochondrial mass and activity is a distinct feature of memory B cell lineage commitment *in vivo*. Using a directed screen of mitochondrial modulators, we identify mitochondrial fission inhibitor, Mdivi-1 as an agent that could enhance mitochondrial mass and function leading to augmented memory B cell differentiation. The enhanced memory B cell responses mediated by Mdivi-1, translated to more robust recall responses upon secondary antigen exposures. Moreover, Mdivi-1 when used in combination with subunit (SARS-CoV2) and inactivated (H1N1 influenza) vaccines led to remarkably improved vaccine efficacies and protection from lethal viral (H1N1) challenge. Single-cell transcriptomics revealed enhanced commitment to memory lineage differentiation in B cells following Mdivi-1 treatment. We propose that mitochondrial modulators such as Mdivi-1 are a novel class of “immune enhancers” that specifically reinforces immunological memory and could be broadly applied to improve the fidelity of immune responses and vaccine efficacies.

## Introduction

Naïve T and B lymphocytes undergo extensive metabolic changes during their differentiation to effector and memory cells, which clear the invading pathogens during primary responses and provide long-term immunity to secondary exposures^1,2^. Though naïve T and B cells are in a state of metabolic quiescence, upon exposure to their cognate antigens, they rapidly alter their metabolic outputs to acquire distinct metabolic states associated with effector and memory phenotypes^1–4^. During the acute phases of clonal expansion and effector differentiation, adaptive immune cells are engaged in anabolic processes and rely on aerobic glycolysis as the primary source of energy^1–3^. Whereas, during memory differentiation, adaptive immune cells switch to catabolic programs which largely rely on mitochondrial oxidative phosphorylation to augment their survival in a nutrient scarce environment^1–3^. Several recent studies have shown that acquisition of distinct metabolic states in effector and memory T cells involves specific structural and morphological changes in the mitochondria^5–8^. The precise mechanisms regulating mitochondrial dynamics and their influence on effector and memory cell fate decisions is an active area of investigation.

The structural and morphological changes in mitochondria are mediated through fission and fusion processes which are proposed to have profound consequences on mitochondrial function^9^. Mitochondrial fission is regulated by the GTPase dynamin related protein-1 (Drp1) which is recruited by adaptor proteins and undergoes assembly into oligomeric spirals to pinch mitochondrial membranes^9–11^. Mitochondrial fusion on the other hand is orchestrated by the fusion of outer and inner mitochondrial membranes by mitofusins (Mfn 1 and Mfn2) and optic atrophy protein 1 (Opa1), respectively^9,12,13^. In effector T cells, increased fission activity promotes generation of smaller, more fragmented mitochondria, while, memory T cells display enhanced fusion activity leading to hyper-fused mitochondrial state with elaborate cristae networks^2,5,6^.

Intriguingly, in cultured T cells a hyper-fused and elongated mitochondrial state imposed by pharmacological inhibition of fission using mitochondrial division inhibitor (Mdivi-1) combined with fusion promoter (M1) leads to attainment of memory-like phenotypes^6^, whereas genetic deletion of mitochondrial fusion protein, Opa1 in T cells restricts survival of memory T cells^6^. How the altered mitochondrial dynamics mediated by fission and fusion processes regulate T and B cell immune responses *in vivo* has not been studied.

Here, we aimed to understand the role of mitochondrial function and the ensuing metabolic changes in B cells undergoing differentiation *in vivo*. Our studies revealed increased mitochondrial mass and function in germinal center (GC) B cells during immune responses and a further enrichment of mitochondrial activity signatures as B cells differentiated into memory lineage. We identified the mitochondrial fission inhibitor, Mdivi-1 as an agent that could enhance mitochondrial mass and activity in GC B cells and further augment memory B cell differentiation. Moreover, Mdivi-1 when used as an immuno-modulator with inactivated (H1N1 influenza) and subunit (SARS-CoV2) vaccines led to remarkably improved vaccine efficacies. Single cell (sc) RNA seq displayed enhanced memory lineage differentiation kinetics in B cells upon Mdivi-1 treatment which translated to superior protection from lethal H1N1 challenge. In summary, we propose that modulators of mitochondrial dynamics, such as Mdivi-1, are a novel class of “immune enhancers” that specifically foster the generation of humoral memory leading to highly persistent immune responses.

## Results

### Modulation of mitochondrial function enhances immunological memory

While extensive changes in mitochondrial structure and function have been reported to accompany T cell differentiation, how mitochondrial dynamics are altered during B cell differentiation are not well-understood. To characterize changes in mitochondrial dynamics during B cell differentiation we immunized C57Bl/6 wild type (WT) mice with 4-Hydroxy-3-nitrophenylacetyl hapten conjugated to chicken gamma globulin (NP-CGG) and examined naïve, Germinal Center (GC) and memory B cell populations at day 21 post-immunization. We observed a progressive increase in mitochondrial mass tracking closely with successive stages of antigen-specific (NP+) B cell differentiation (Naïve<GC<Memory) within the same mice (**Fig. 1a**). We also measured fatty acid oxidation which is posited to be the primary pathway feeding into mitochondrial oxidative phosphorylation in GC B cells^14^, by monitoring the oxygen consumption rate (OCR) in differentiating B cells following administration of long-chain fatty acid (BSA-palmitate) 40 mins before harvesting splenic B cells at day 15 post-immunization with sheep red blood cells. Notably, our studies revealed significantly higher OCR in memory B cells compared with naïve and GC B cells (**Fig. 1b**), together showing, that B cells undergoing differentiation to memory lineage exhibit enhanced mitochondrial mass and activity (OCR).

**Figure 1.**
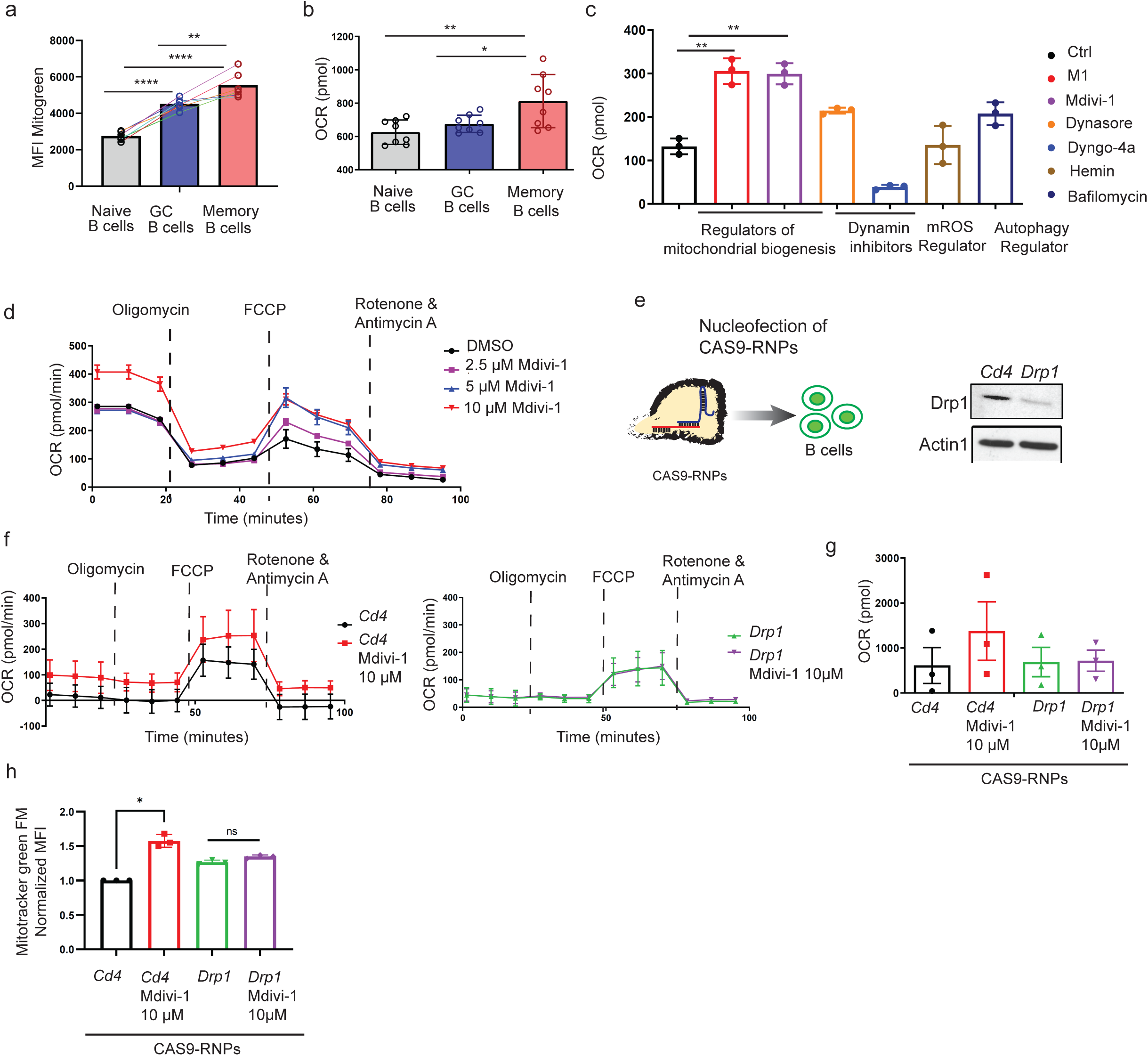
Mitochondria fission inhibitor, Mdivi-1, enhances oxygen consumption rate of stimulated B cells. a) Mice (n= 6) were immunized with NP-CGG conjugated with alum. Mice were sacrificed on Day 21 post immunization and spleen were harvested. Displayed is the bar graph of median fluorescence intensity (MFI) of MitoTracker Green FM in naïve, GC and memory B cells, each dot represents individual mice. b) Mice (n=9) mice were immunized with sheep red blood cells (SRBC) on Day 0 and Day 4. On Day 15 post-immunization mice were injected with BSA conjugated with palmitate 40 mins before sacrificing mice. Naïve, GC, Memory B cells were MACSs, cells per well were plated in eight well per sample and seahorse assay was performed. Displayed is a bar graph for basal oxygen consumption rate (OCR) in respective B cells. c) Naïve splenic B cells were isolated from C57/B6 mice, B cells were next stimulated with CD40 and anti-IgM for 24 hours. After 24 hours molecules were added M1 fusion promoter (20µM), Mdivi-1 (10µM), Dynasore (10µM), Dyngo-4a (10µM), Hemin (60µM) and Bafilomycin (10nM). At 48 hours, seahorse was performed to determine the oxygen consumption rate, cells were plated in triplicates. DMSO treated stimulated cells were used as control. Displayed is a bar graph of basal OCR in each condition. d) Naïve B cells were stimulated with CD40 and anti-IgM for 24 hours. Mdivi-1 was added to cells at the concentration of 2.5 µM, 5 µM and 10 µM and seahorse assay was performed. Displayed is the time lapse oxygen consumption rate in different conditions. e) Naïve B cells were electroporated with Drp-1 CAS9-Ribonucleoproteins (RNPs), followed by stimulation with CD40 and anti-mouse IgM for 48 hours, treated with or without Mdivi-1 (10µM). Cd4 CAS9-RNPs and DMSO were used as control. f) Seahorse was performed to determine the oxygen consumption rate, cells were plated in triplicates. g) Graph represents total OCR in each condition from f). h) Graph represents quantification of mitochondrial mass in respective condition. Statistical significance was calculated using one-way ANOVA for a-c; in prism p=0.01*, p=0.005**, p=0.0001***, p<0.0001****.

With these observations in mind, we asked if we could influence memory B cell differentiation by altering mitochondrial structure and activity. To this end, we designed a directed screen with small molecule regulators of mitochondrial structure and function to identify pharmacological agents that could be used to simulate mitochondrial mass and activity profiles observed in memory B cells. We examined OCR in B cells upon treatment with four known classes of mitochondrial modulators, including those- 1) regulating mitochondrial biogenesis by altering fusion (M1)^15^ and fission processes (Mdivi-1)^16^; 2) affecting mitochondrial localization through inhibition of dynamin GTPase activity (Dynasore and Dyngo-4a)^17,18^; 3) impacting mitochondrial ROS levels (Hemin)^19^ and; 4) influencing mitophagy (Bafilomycin)^20^. Treatment of B cells in culture with these four classes of mitochondrial modulators identified, mitochondrial division inhibitor-1 (Mdivi-1) as one of the agents that could enhance total OCR in B cells when compared with vehicle only control (**Fig. 1c**). Though other regulators of mitochondrial biogenesis such as the fusion inducer, M1 also supported increased OCR, we focused on Mdivi-1 for further studies given its better characterized mechanism of action. Mdivi-1 is a potent inhibitor of dynamin-related protein 1 (Drp1) which is a critical effector molecule for mitochondrial fission process^9–11^. Treatment of stimulated B cells with Mdivi-1 led to a dose-dependent enhancement of mitochondrial mass and function characterized by an increase in total as well as maximal respiration, while Cas9-Ribonucleoprotein (Cas9-RNPs) mediated targeting of Mdivi-1 target, Drp1, negated these effects (**Fig. 1d-h**).

We then performed functional assays to test the effects of Mdivi-1 treatment towards augmenting mitochondrial mass and activity as well as memory B cell differentiation *in vivo*. Treatment with Mdivi-1 in mice undergoing active immune responses (**Fig. 2a**), led to an overall increase in mitochondrial mass in activated germinal center B cells (**Fig. 2b** **and 2c and Sup Fig. 1e**). Moreover, germinal center (**Fig. 2d**) and IgG1 expressing memory B cells (**Fig. 2e**) and also displayed enhanced mitochondrial activity as quantified by CMXROS, a probe which specifically accumulates in a mitochondrial membrane potential (ᴪ_m_) dependent manner. Importantly, treatment with Mdivi-1 led to an ∼2.5-fold increase in the absolute numbers of IgG1 memory B cells (**Fig. 2f** **and Sup Fig. 1a**), clearly demonstrating that enhanced mitochondrial activity during an ongoing immune response can foster memory B cell differentiation. We further validated the role of Mdivi-1 towards enhancing memory B cell differentiation by assessing antigen-specific recall responses. For these studies, mice initially immunized with NP-CGG in Alum were re-challenged after 2 months with a sub-optimal regimen of antigen, NP-CGG in PBS and high affinity antibody responses were measured by ELISA. All mice (100%) immunized with NP-CGG and treated with Mdivi-1 during primary response responded upon re-challenge, compared to a bi-modal distribution of the response (50% high responders and 50% low responders) observed in cohort of mice treated with vehicle alone during primary immunizations (**Sup Fig. 1b-d**).

**Figure 2.**
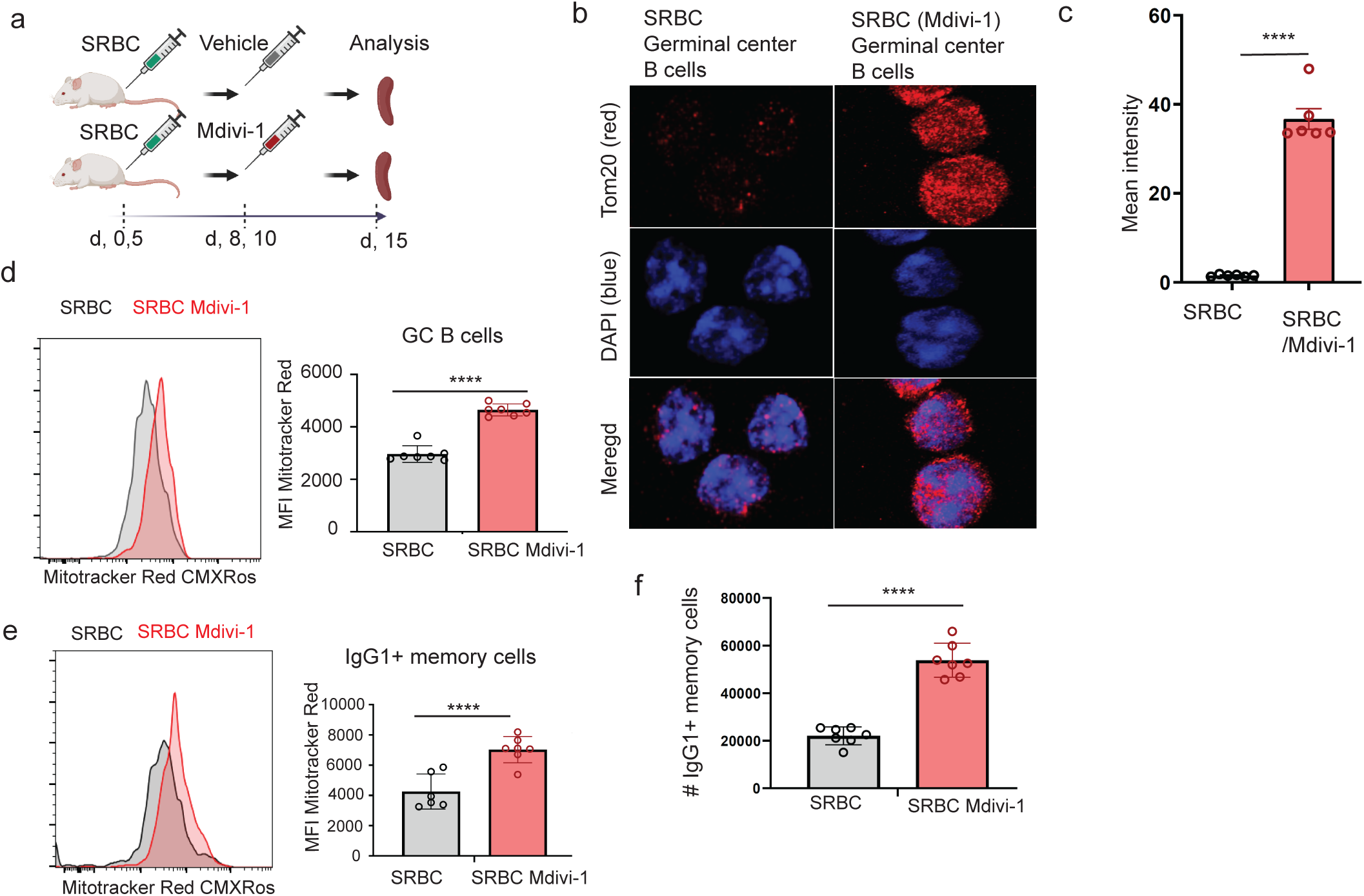
Inhibition of mitochondria fission during immunization enhances mitochondrial mass and number of memory B cells. a) Experimental schematic, mice (n=7) each in group were immunized with SRBC on Day 0 and Day 5. Mice in each group on Day 8 and Day 10 were given Vehicle or Mdivi-1 at 2.5mg/kg. b) Isolated GC B cells were stained with Tom20 (red) to stain mitochondria and DAPI to stain nucleus. Cells were imaged using confocal microscope. Images are of GC B cells from mice immunized with SRBC with vehicle and SRBC with Mdivi-1. c) Mean intensity of each red dot, i.e. mitochondria stained with Tom20 in displayed images in b), using ImageJ software. d) Histogram and graph of mitochondrial mass measured by Mitotracker red MFI in GC B cells and e) Memory B cells in vehicle control and Mdivi-1 treated mice group. f) Displayed graph represents absolute number of CD38+ IgG1+ memory B cells in SRBC with vehicle and SRBC with Mdivi-1. Each dot represents single mouse in the group. Statistical significance was calculated using t-test for c-f; in prism p<0.0001****.

### Pharmacological inhibition of mitochondrial fission improves the efficacy of influenza vaccine

Since immunological memory is one of the core principles for effective vaccines, we next evaluated the effect of Mdivi-1 on vaccine efficacy in a murine model of H1N1 influenza infection. For these studies we used four different cohorts of mice: 1) unvaccinated, 2) treated with Mdivi-1 alone, 3) vaccinated (inactivated H1N1 virus) and, 4) vaccinated in combination with Mdivi-1 (Vaccine /Mdivi-1). 60 days post-vaccination all four groups were exposed to a lethal dose of H1N1 virus (**Fig. 3a**). As expected, mice with little or no protection against H1N1 infection appeared feeble and showed rapid mortality in unvaccinated (100%) and Mdivi-1 only (80%) cohorts (**Fig. 3b** **and Sup Video. 1 a-d**). Vaccinated mice also exhibited limited protection (40% survival) to a lethal H1N1 challenge. However, rather strikingly, the Vaccine/Mdivi-1 cohort showed vigorous health and 90% overall protection to a lethal H1N1 challenge compared to the other cohorts (**Fig. 3b** **and Sup Video. 1 a-d**). Though most animals in the unvaccinated, Mdivi-1 treated and vaccinated groups could not survive the viral challenge beyond day 8, the Vaccine/Mdivi-1 treated group began to show signs of recovery at day 4 post-infection, as reflected by the significantly less weight-loss when compared with unvaccinated, Mdivi-1 alone and vaccinated group, at day 6 post-infection (**Fig. 3c** **and Sup. Fig. 2a)**. Histopathological analysis of mice that succumbed to the infection (only one mouse in the Vaccine /Mdivi-1 group) also presented less severe lung disease in Vaccine/Mdivi-1 cohort compared with other cohorts (**Sup. Fig. 2b**). Together, these results demonstrate that small molecule enhancers of mitochondrial function *in vivo* could represent a novel strategy to boost vaccine efficacies.

**Figure 3.**
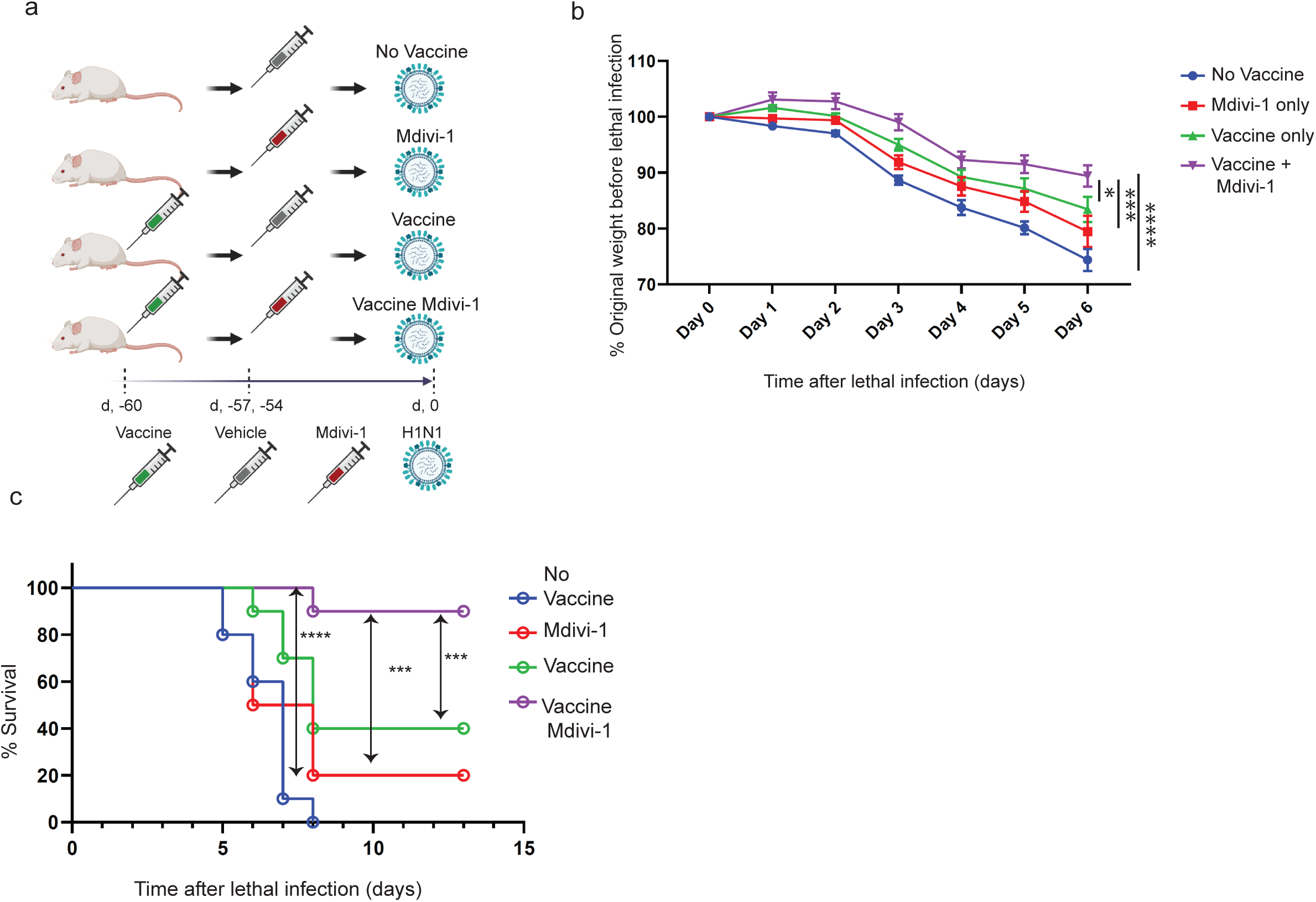
Mdivi-1 enhances flu vaccine efficacy in influenza mouse model. a) Schematic of experimental design to test the effect of Mdivi-1 on H1N1 vaccine efficacy. b) Graph represents normalized weights for each mouse in respective groups post lethal H1N1 infection challenge. c) Kaplan-Meier event free survival curve of mice from respective groups lethally challenged with H1N1 virus. Mice were sacrificed if the weight reached 70% of the original weight or appeared very sick. N=10 per group. Vehicle only was used as control. Statistical significance was calculated using 2-way ANOVA and corrected for multiple comparisons for b), and log-rank test for c); in prism. p=0.04*, p=0.005**, p=0.0001***, p<0.0001****.

### Inhibition of mitochondrial fission augments memory B cell differentiation following H1N1 vaccination

Mdivi-1 treatment following vaccination was associated with strong protection against lethal H1N1 challenge, therefore, to characterize the changes in the immune cell subsets affected by this treatment and understand the underlying mechanism, we performed a 5’ directed single-cell RNA (sc-RNA) sequencing. We examined the mediastinal lymph nodes (mLNs), the draining lymphoid tissues for lungs, to capture the early kinetics of the immune response following a) vaccination (at day 10) and b) subsequent viral infection (at day 34 post-vaccination or day 4 post-viral challenge) (**Fig. 4a**) in mice vaccinated with or without Mdivi-1. sc-RNA sequencing revealed profound changes in B and T cell differentiation upon vaccination and H1N1 viral challenge (**Fig. 4b**). Given the impact of Mdivi-1 treatment on B cell differentiation, we first focused on characterizing changes in B cell dynamics. Our analysis highlighted strong activation signatures (high *CD69* and *CD83* expression) in B cells within mLNs with or without Mdivi-1 treatment at both surveyed time-points (**sup Fig. 3a &b**). Initial uniform manifold approximation and projection (UMAP) clustering separated bulk B cells into populations with an activated phenotype either harboring diverse Immunoglobulin (Ig) light chain repertoires (**Fig. 4b**; pre-dominant cluster in circle) or smaller clusters expressing clonally distinct Ig light chains (**Fig. 4b**). However, no apparent differences were observed in the gene expression profiles between the pre-dominant B cell cluster and the smaller clonally distinct clusters, besides the Ig light chain gene segments being the only major differentially expressed genes between the clusters. Moreover, the distribution of the pre-dominant or comparatively minor, clonally distinct clusters were neither altered based on the time-point nor based on the treatment group. Hence, we performed two iterative rounds of UMAP clustering to further dissect the pre-dominant B cell cluster using specific markers for B cell differentiation (*CD38, CD83, Ccr6, Bach2, Hhex, Irf4*) into subsets of activated, early GC, precursor memory, early memory and mature B cells (**Fig. 4b**; Reclustered B cells, **sup Fig. 3b**).

**Figure 4.**
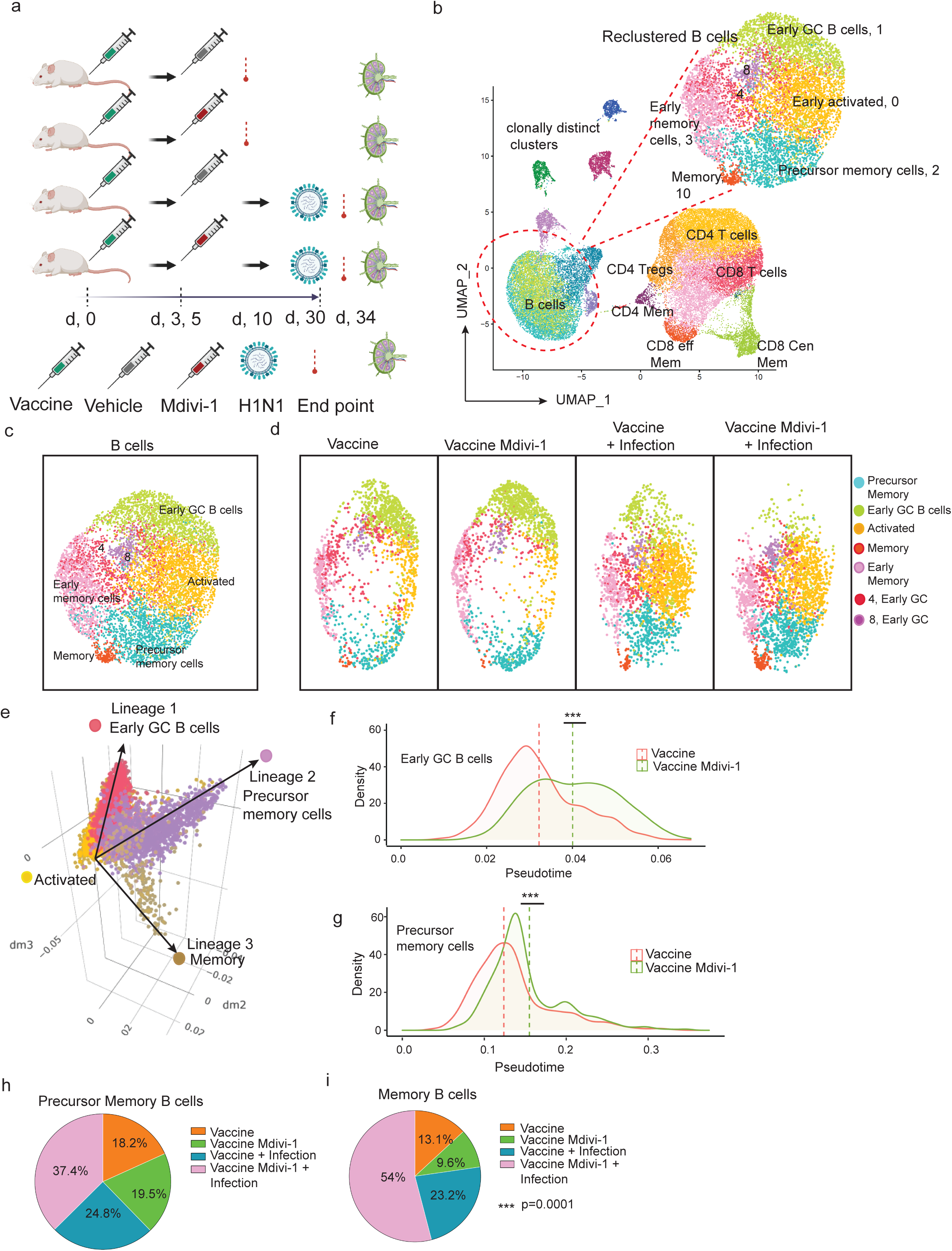
Mdivi-1 reinforces memory B cells differentiation *in vivo* to enhances flu vaccine efficacy. a) Schematic for single cell sequencing of mediastinal lymph nodes from mice (n=5) that were pooled together per group. Vehicle only was used as control. b) UMAPs identifying different immune cells clusters. B cells further re-clustered to pars out distinct subsets, highlighted in the insert. c) UMAPs combined B cells clusters from all the samples. d) UMAPs identifying B cell clusters in mice from different groups. e) Displayed is the pseudo-time lineage maps from B cells depicting activated cells differentiating into lineage 1, Early GC B cells, lineage 2, Precursor Memory cells, lineage 3, Memory. Histograms (x-axis as pseudo-time and y-axis as normalized cell counts) depicting activated cells differentiation to f) lineage 1, early GC B cells; g) lineage 2, Precursor Memory B cells in Vaccine and Vaccine (Mdivi-1) treated mice. h & i) displayed are the pi-charts showing memory B cells distribution in mice from Vaccine, Vaccine (Mdivi-1), Vaccine + Infection and Vaccine (Mdivi-1) + Infection groups. Statistical significance was calculated using Wilcoxon signed rank test for f-g, chi-square test for h-i, p=0.04*, p=0.005**, p=0.0001***, p<0.0001****.

Clusters of early GC (clusters 1,4 and 8; *CD83^++^ CD38^lo^, Irf4^+^*), early memory (cluster 3; *CD38^++^*) and precursor memory (cluster 2; *CD38^++^, Ccr6^+^, Hhex^+^*) B cells were present at day 10 post-vaccination, while, clusters defining the activated (cluster 0; *CD83^++^ CD38^+^*) and mature memory (cluster 10; *CD38^+++^ Bach2^+^*) B cells were more prominent at day 4 post-infection (**Fig. 4c, 4d** and **Sup Fig. 3b**). To further tease out the B cell kinetics we inferred the *in vivo* differentiation trajectories using pseudo-time analysis^21^. We identified three distinct differentiation trajectories culminating into lineage 1: early GC, lineage 2: precursor memory and lineage 3: mature memory B cells (**Fig. 4e**). Intriguingly, pseudo-time analysis revealed a marked acceleration of B cells in their progression through the early GC and precursor memory differentiation trajectories in mice treated with Mdivi-1 when compared with vehicle only controls (**Fig. 4f** **and 4g**). These early changes also translated to a noticeable increase in the frequencies of precursor memory, mature memory B cell and early GC B cell differentiation trajectory, and mature B cells in lungs post-infection with H1N1 in mice vaccinated in combination with Mdivi-1 compared with vaccine only group (**Fig. 4h, 4i, Sup Fig. 3c-e**). Furthermore, using SRBCs and NP-CGG as model antigens we confirmed the enrichment of precursor memory B cell pools and replenishment of fully-differentiated memory cells upon recall responses following treatment with Mdivi-1^22,23^ (**Sup Fig. 4 a-d**), respectively. Together, these findings highlight early effects of Mdivi-1 in enhancing B cell differentiation towards memory lineage.

### Slight changes in T cell differentiation upon Mdivi-1 treatment

Efficient protection against H1N1 infection requires both B and T cell components of the adaptive immune response^24,25^. sc-RNA-seq analysis identified clusters of activated and memory-like CD4 and CD8 T cells in the mLNs (**Sup Fig. 5a and Sup Fig. 5a & b**), and we also examined the changes in T cell differentiation upon H1N1 vaccination with or without Mdivi-1 treatment and the subsequent viral challenge. Based on the expression of known markers and transcriptional signatures from ENCODE database, we were able to define clusters of activated CD4 and CD8 T cells, CD4 effector memory cells, CD8 effector and central memory cells and a minor population of CD4 Tregs (**Sup Fig. 5a**). Pseudo-time analysis and frequency distribution profiles did not show significant differences in the T cell differentiation kinetics in mice treated with or without Mdivi-1 post-vaccination (**Sup Fig. 5b-e**). However, the differentiation towards CD8 effector and central memory subsets were significantly enhanced in Mdivi-1 treated group following the viral challenge (**Sup Fig. 5f and g**). We noticed a strong depletion of T cell subsets from the mLN following viral challenge, suggesting egress of T cells to the effector site (**Sup Fig. 5h**) and indeed, immunohistochemistry readily identified infiltrating CD4 and CD8 cells in lung tissues post-H1N1 infection (**Sup Fig. 5i**). Mice which received vaccination in combination with Mdivi-1 treatment showed significantly higher CD4 and CD8 infiltration post H1N1 infection (**Sup Fig. 5i and j**), consistent with the enhanced T cell differentiation kinetics observed by sc-RNA-seq analysis in mLN post viral infection (**Sup Fig. 5f and g**). Together, these results suggest that Mdivi-1 treatment with vaccination, atleast at the administered dose and the time-points surveyed, does not have a major impact on T cell differentiation, and the changes in T cell differentiation observed upon viral challenge in Mdivi-1 treated groups could likely be an indirect consequence of the effect of Mdivi-1 treatment on other (immune) cell types.

### Mdivi-1 treatment promotes differentiation to lineages associated with enhanced mitochondrial function

We observed increased mitochondrial mass and activity (OCR and ᴪ_m_) in differentiating B cells upon Mdivi-1 treatment (**Fig. 2b-d**). To further illuminate the underlying mechanism, we examined the enrichment of transcriptional signatures associated with mitochondrial function in distinct B cell lineages following Mdivi-1 treatment. Compared to the early GC B cells and fully-differentiated memory B cells, the precursor memory cells displayed a strong enrichment in gene signatures linked with mitochondrial function such as, oxidative phosphorylation, electron transport chain, beta-oxidation and glycolysis (**Fig. 5a**). Moreover, in addition to monitoring the mitochondrial activity profiles in specific B cell clusters, we also assessed the transcriptional signatures associated with mitochondrial function in actively differentiating lineages. To this end, we compared the enrichment of signatures associated with mitochondrial function in B cells at early (start) versus late (end) stages of differentiation within the precursor memory and mature memory lineage trajectories (**Fig. 5b**). Interestingly, B cells making their way down the precursor memory differentiation path showed a significant enrichment in oxidative phosphorylation, beta-oxidation and glycolytic gene signatures at late compared to early stages of pseudo-time lineage trajectories (**Fig. 5c**). Whereas, B cells adopting the mature memory B cell lineage path showed a significant downregulation of mitochondrial activity signatures at late compared to early stages of differentiation (**Supplementary Fig. 6a**). Taken together, these data imply that Mdivi-1 treatment and the ensuing changes in mitochondrial mass and function potentially skews B cell differentiation towards precursor memory lineage that are inherently associated with enhanced mitochondrial activity.

**Figure 5.**
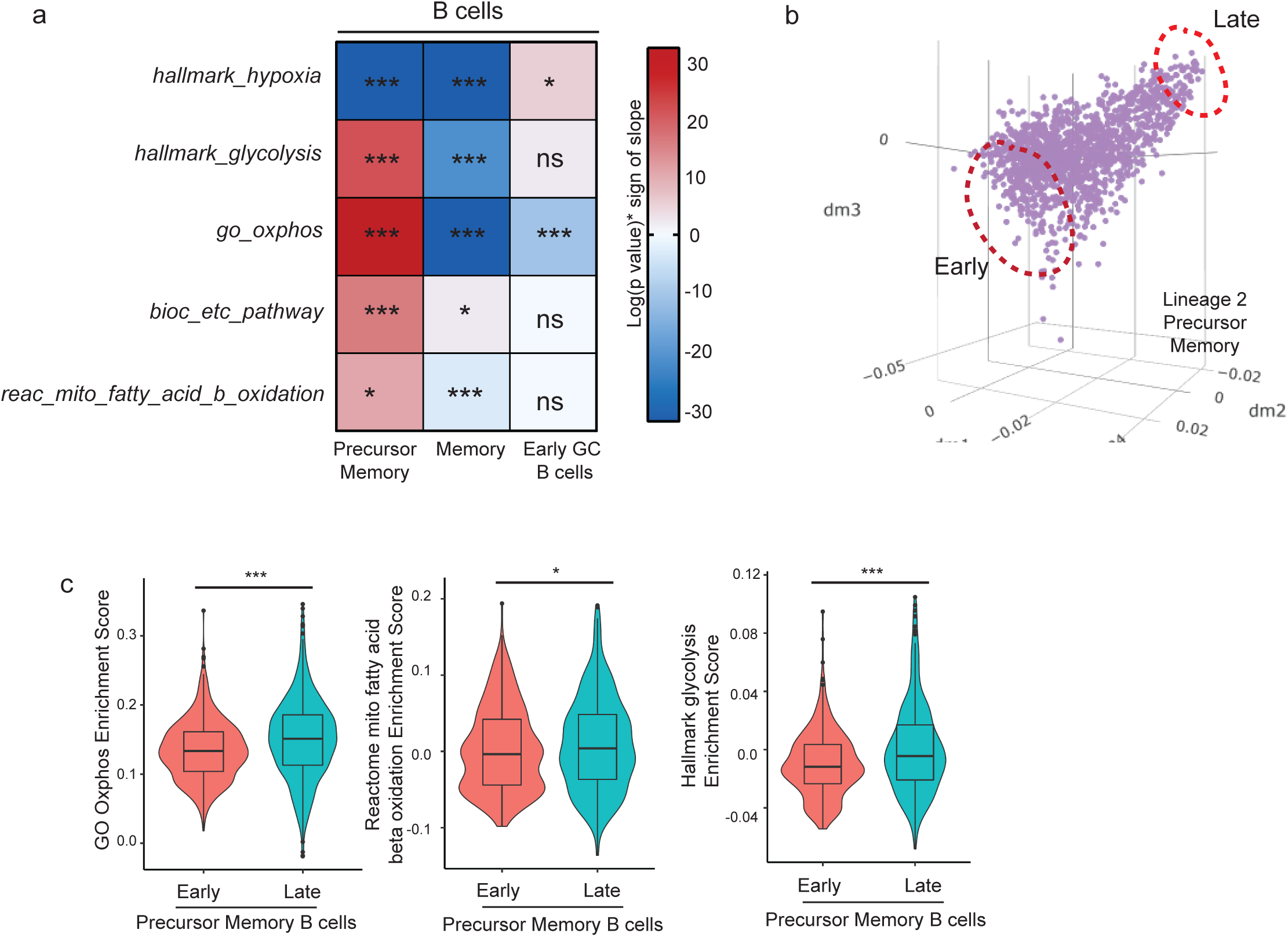
Precursor Memory B cells show strong enrichment of pathways associated with mitochondrial function. a) Normalized enrichment score of genes associated with metabolic pathways. b) top and bottom 10% of grouped B cells by pseudo-time and labelled the groups as “early” (start of the lineage 2) and “late” (end of the lineage 2). c) Normalized enrichment score of GO oxphos (left), Mitochondrial Fatty acid beta oxidation (center) and Hallmark glycolysis (right) pathways of the top and bottom 10% of grouped B cells in b). Statistical significance was calculated using linear model for K, p=0.04*, p=0.005**, p=0.0001***, p<0.0001****.

### Inhibition of mitochondrial fission as a broad strategy to boost vaccine efficacies

Motivated by the strong effects of Mdivi-1 towards increasing the potency of an inactivated viral vaccine, we evaluated the broad applicability of this strategy to boost vaccine efficacies. We tested the activity of Mdivi-1 in combination with a surrogate SARS-CoV2 subunit vaccine. We used the receptor binding domain of the S1 subunit (S1-RBD) of the SARS-CoV2 spike glycoprotein and immunized mice with recombinant S1-RBD conjugated to alum followed by treatment with vehicle or Mdivi-1 (**Fig. 6a**). We observed a significant increase in the total number of antigen (S1-RBD)-specific B cells in mice immunized with alum-conjugated S1-RBD compared with mice immunized with alum alone or alum in combination with Mdivi-1 (**Fig. 6b**). We then measured the robustness of memory/recall response by re-challenging all 4 groups of mice with S1-RBD without alum 30 days post-initial immunization by monitoring serum antibody titers at day 7 post re-challenge. While mice immunized with alum alone or alum with Mdivi-1 showed no detectable levels of S1-RBD specific antibodies, the group of mice initially immunized with S1-RBD and treated with Mdivi-1 showed significantly higher (∼1.5 fold) levels of S1-RBD specific antibodies compared with vehicle only group (**Fig. 6c**). Moreover, mice treated with Mdivi-1 showed higher frequency of IgG1 positive B cells in spleen compared with vehicle treated group (**Fig. 6d**). Intriguingly, the antibodies from the sera of S1-RBD immunized and Mdivi-1 treated mice were ten-folds better in their ability to neutralize the infection by SARS-CoV2 spike glycoprotein expressing pseudoviral particles in Vero-E6 cells, compared to sera isolated from S1-RBD with vehicle only group (**Fig. 6e**). We also compared the neutralization activity towards the D614G mutant spike glycoprotein^26^ expressing pseudoviral particles and observed significantly higher neutralization in sera from immunized mice treated with Mdivi-1 compared with vehicle only controls (**Fig. 6f**). Collectively, these studies describe that modulators of mitochondrial dynamics such as Mdivi-1 could be used as a novel class of “immune enhancers” broadly applicable to different classes of vaccines.

**Figure 6.**
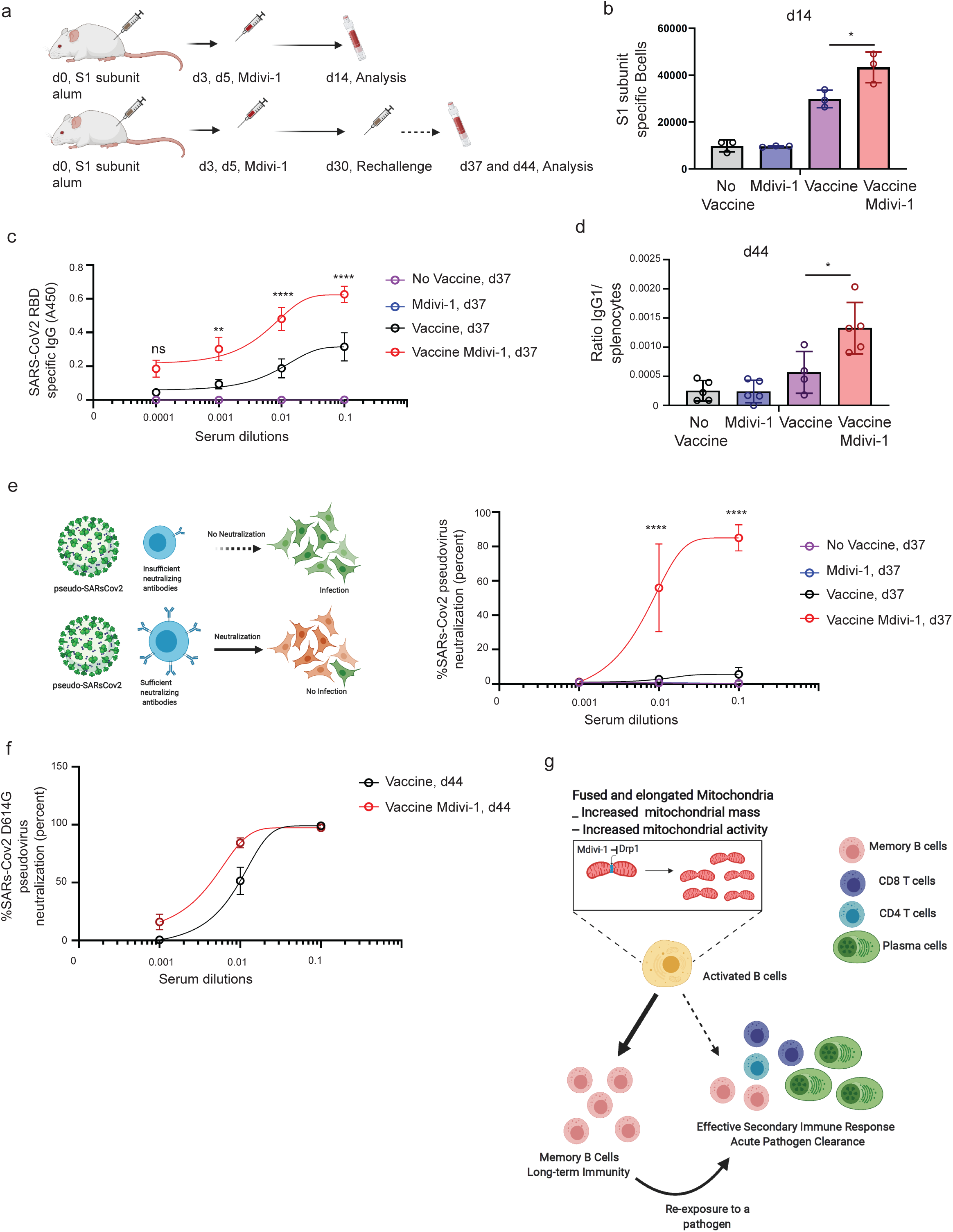
Inhibition of mitochondrial fission leads to more potent immune responses against SARS-COV2 spike protein. a) Schematic for studying B responses to surrogate SARS-COV2 vaccine. Mice (n= 8), were immunized with S1 subunit of coronavirus spike protein conjugated with alum on Day 0. Day 3 and Day 5 post immunization mice were given Mdivi-1 (2.5mg/kg). Unimmunized mice, mice only treated with Mdivi-1 and Immunized mice treated with Vehicle were used as controls. Spleens from mice (n=3) each in respective groups were harvested and stained for B cells specific to S1 subunit, b) Graph represents the S1 subunit specific B cells measured by flow cytometry in spleens from mice immunized with S1 subunit with or with Mdivi-1 and control mice. c) Antibody titers in serum collect on Day 37 against S1 subunit measured by ELISA from mice n=5 each in group. d) Ratio of IgG1 positive cells in spleen measured by flow cytometry analysis, d44. e) ACE-2 receptor expressing cells were infected with SARS-Cov2 pseudovirus particle in presence of sera isolated from mice on d37, sigmodal dose curve of percent neutralization of e) wild type SARS-Cov2 pseudovirus f) D614G mutant SARS-Cov2. g) Proposed Model on how alteration in mitochondrial dynamics could lead to long-term immunity. Statistical significance was calculated using one-way Anova for b and d, two-way anova for c, e and f, in prism p=0.01*, p=0.005**, p=0.0001***, p<0.0001****.

## Discussion

Our studies highlight enhanced mitochondrial mass and activity as an intrinsic feature of memory B cells which prompted us to devise a pharmacological strategy that could boost the efficacy and robustness of immune responses through augmenting memory B cell differentiation (**Fig 6f**). Using a directed screen, we identified Drp1 antagonist, Mdivi-1 as an agent that can enhance mitochondrial function while promoting the generation of memory B cells *in vivo* (**Fig 6f**). Importantly, Mdivi-1 treatment when combined with model immunogens and experimental vaccines had profound effects on improving the overall efficacy of immune responses and providing protection against viral exposure (**Fig 6f**). In view of our rapidly evolving knowledge regarding the metabolic regulation of immune responses, these studies provide an important proof-of-concept, rationalizing the design and use of metabolic modulators such as, Mdivi-1 to enhance the fidelity of immune responses.

Extensive mitochondrial remodeling during T and B cell activation have been previously reported^5–7,27,28^, but whether the changes in mitochondrial dynamics play an active role during acquisition of specific cell fates in differentiating T and B cells remains unclear. In this regard, we show that transient alterations in mitochondrial mass and activity could have conspicuous effects on cell fate decisions, particularly towards memory lineage differentiation in B cells. Studies in cultured T cells, where mitochondrial dynamics during differentiation are better characterized, have shown that pharmacological induction of mitochondrial fusion by a combination of Mdivi-1 and fusion inducer, M1, promotes memory-like cellular phenotypes, while genetic ablation of Opa1, which mediates mitochondrial fusion was shown to be detrimental to memory fate^6^. Mechanistically, the elongated mitochondrial morphology and the elaborate cristae network resulting from mitochondrial fission inhibition, are postulated to support catabolic metabolism required for memory cell survival through more efficient mitochondrial respiration and function^2,5,7^. Consistent with these notions, our studies reveal that transient Mdivi-1 treatment *in vivo* during an ongoing immune response enhances B cell differentiation towards a precursor memory fate which are enriched for higher mitochondrial activity signatures. It is noteworthy, while the precursor memory cell pools showed an enrichment of mitochondrial activity signatures, the fully differentiated memory B cells did not, suggesting that the mitochondrial activity is perhaps transiently enhanced as B cells differentiate towards memory lineage. Our findings are in concordance with recent studies describing essential roles of mitochondrial oxphos signatures in differentiating GC B cells^14,29^. Understanding the precise molecular mechanism through which mitochondrial remodeling activity of Mdivi-1 leads to such striking and lasting changes in B cell differentiation would need further investigation.

The dose of Mdivi-1 administered in our studies are ∼20 times lower than doses shown to be well-tolerated in rodents^30^ and ∼5 to 10 times lower than doses reported to promote anti-proliferative effects in tumor studies^30^. Moreover, we wanted to transiently target immune cell populations that recently underwent activation in response to their cognate antigens but have not yet committed to specific lineages and therefore, we rationally timed the delivery of Mdivi-1 within the first few days post-immunization/vaccination. Interestingly, under these dosing regimens we observed clear changes mediated by Mdivi-1 treatment on B cell differentiation *in vivo*, however the immediate effects of Mdivi-1 in T cell differentiation, at least in the H1N1 model, were not easily noticeable. These observations suggest that B cells undergoing differentiation are more sensitive to Mdivi-1 induced mitochondrial remodeling. Nonetheless, we did observe significant changes in T cell differentiation, particularly in CD8 T cells during recall responses which indicates, albeit indirectly, Mdivi-1 treatment can influence T cell differentiation. Of note, it would be interesting to investigate how differentiating B cell in presence of Mdivi-1 could influence CD8 T cell differentiation in these contexts, as has been reported previously^31–33^.

Given the positive effects of Mdivi-1 to reinforce the generation of immune memory, our studies clearly show that our approach could be applied as a general strategy to boost vaccine efficacies. We present data demonstrating strong effects of Mdivi-1 on enhancing the efficacy of two different vaccine types, 1) an inactivated H1N1 viral vaccine and 2) a spike glycoprotein subunit vaccine against SARS-CoV2. Mdivi-1 treatment promoted more persistent and qualitatively superior immune responses as demonstrated by protection against lethal H1N1 challenge and the neutralization of wildtype and mutant spike glycoprotein expressing pseudoviral particles. While not formally tested here, our studies suggest that the enhanced memory B cell differentiation upon Mdivi-1 treatment could also allow for secondary diversification of B cell responses, thus equipping the immune system to better counteract antigenic drifts commonly associated with pathogen evolutions. In summary, we show that inhibition of mitochondrial fission reinforces B cell immunological memory leading to improved immune responses and vaccine efficacies.

## Acknowledgements

We thank Dr. Sumit Chanda and members of the Chanda lab (Scripps Research, La Jolla, CA) for providing us Vero cells and SARS-CoV2 pseudoviral particles. We also thank SBP animal facility, histology core and Genewiz for performing single cell RNA seq. A.B. was supported by NIH/NCI (1R01CA200643-01A1), and Institutional startup funds. This study was funded by National Health Institute grant R01AI122344, SBP Institutional startup to A.B. and Styx Biotechnologies Incorporated.

## Author Contributions Statement

A.S., V.S., A.T. and A.B. designed the studies, analyzed the data and wrote the manuscript. R.R. and M.D.T. provided suggestions for the B cell and RNA-Seq studies, respectively. L.H. and O.S. performed the bioinformatics analysis of sc RNA-Seq data. B.H. assisted with the animal studies. All authors reviewed and edited the manuscript.

## Competing Interests Statement

A.B, A.T., A.S and V.S are co-founders of Styx Biotechnologies Incorporated, San Diego, CA.

## Data Availability

All genome-wide sequencing datasets have been deposited to Gene Expression Omnibus (GEO) repository, accession number GSE201082. Any data and reagents will also be made available upon request.

---

**Supplementary Figure 1.**
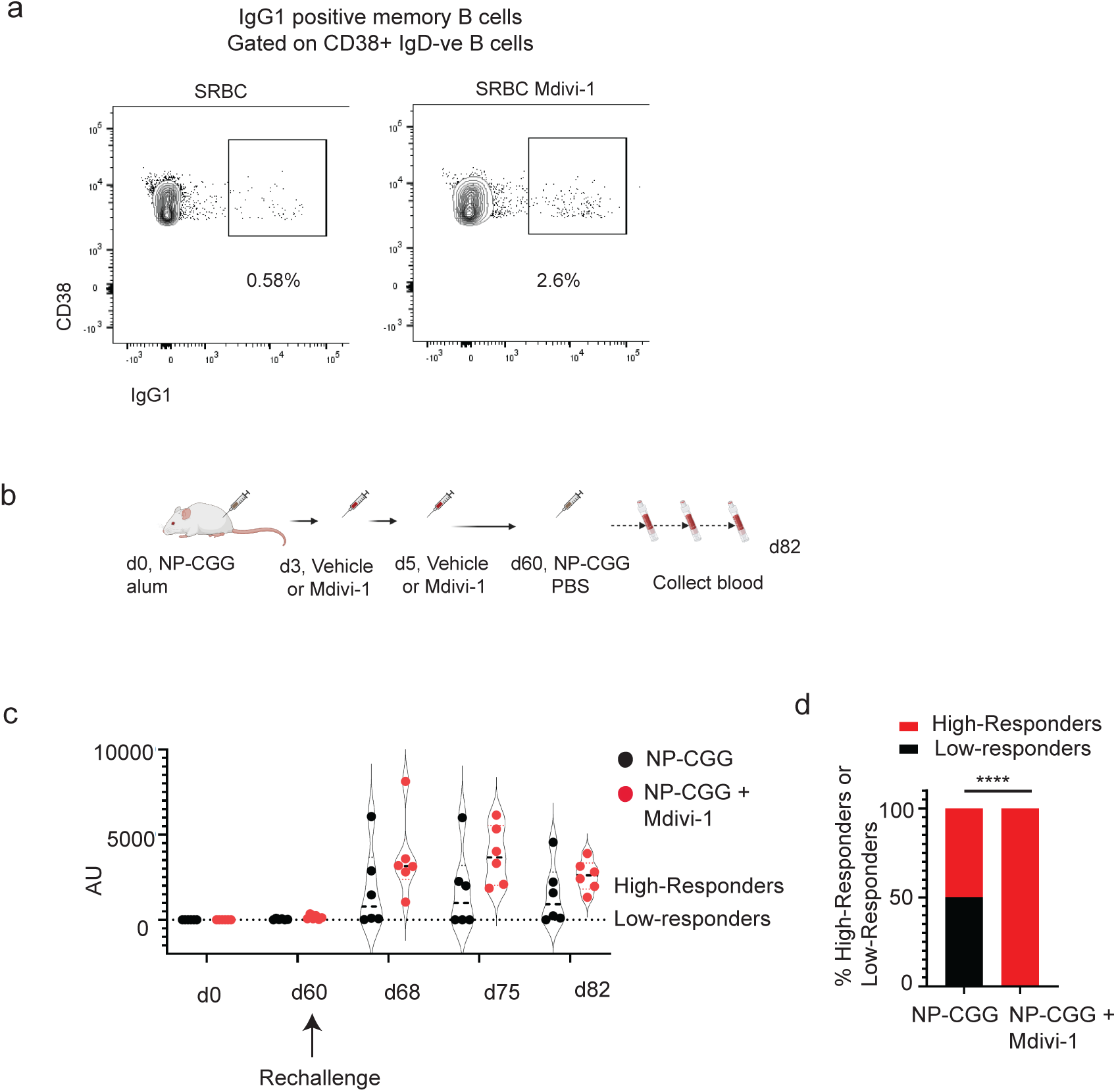
Modulation of mitochondrial function during immunization enhances immune memory. a) Gating strategy to identify IgG1 positive memory B cells. b) Experimental design for studying NP-CGG immunization along with or without Mdivi-1 treatment. c) Violin plots showing the NP-2 specific antibody production measured by ELISA in control and Mdivi-1 treated mice upon re-challenge with NP-CGG in PBS on Day 61. Mice were bled on Days 0, 60, 68, 75 and 82. d) Displayed is the contingency responder’s graph quantified as high or low responders upon antigen re-challenge (chi-square test). Statistical significance was calculated using; chi-square test for d; in prism p<0.0001****.

**Supplementary Videos.**
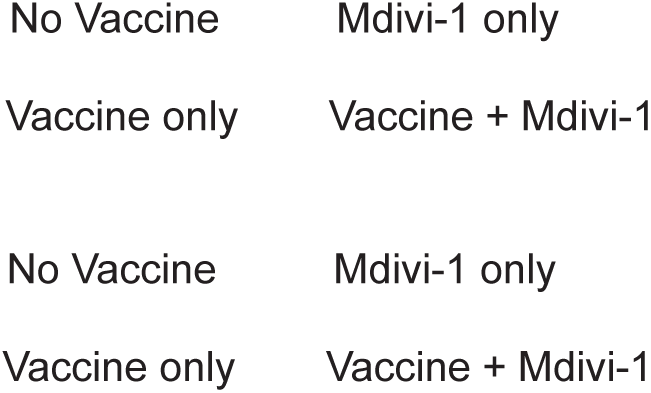
Vaccine with Mdivi-1 delays the flu disease progression in mice. Video links of single mice or groups of mice from No vaccine, Mdivi-1 only, Vaccine only, and Vaccine + Mdivi-1 treated cohorts.

**Supplementary Figure 2.**
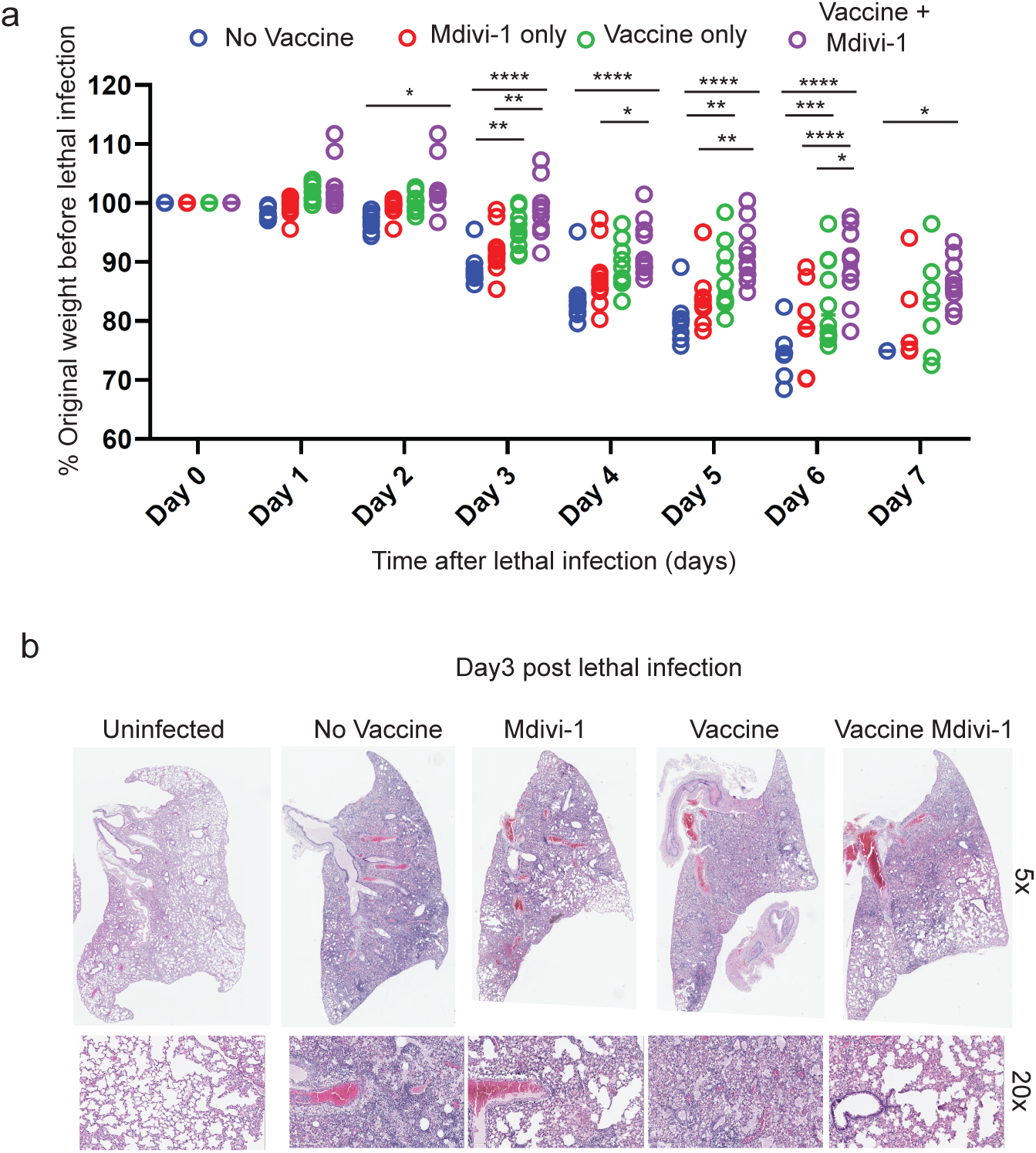
Vaccine with Mdivi-1 delays the flu disease progression in mice. a) Graph represents normalized weights for each mouse in respective groups (n=10) post lethal H1N1 infection challenge. b) H&E staining of lungs (left lobe) at 5x and 20x from uninfected and mice infected with H1N1 from respective groups. Statistical significance was calculated using 2-way ANOVA and corrected for multiple comparisons for a); in prism. p=0.04*, p=0.005**, p=0.0001***, p<0.0001****.

**Supplementary Figure 3.**
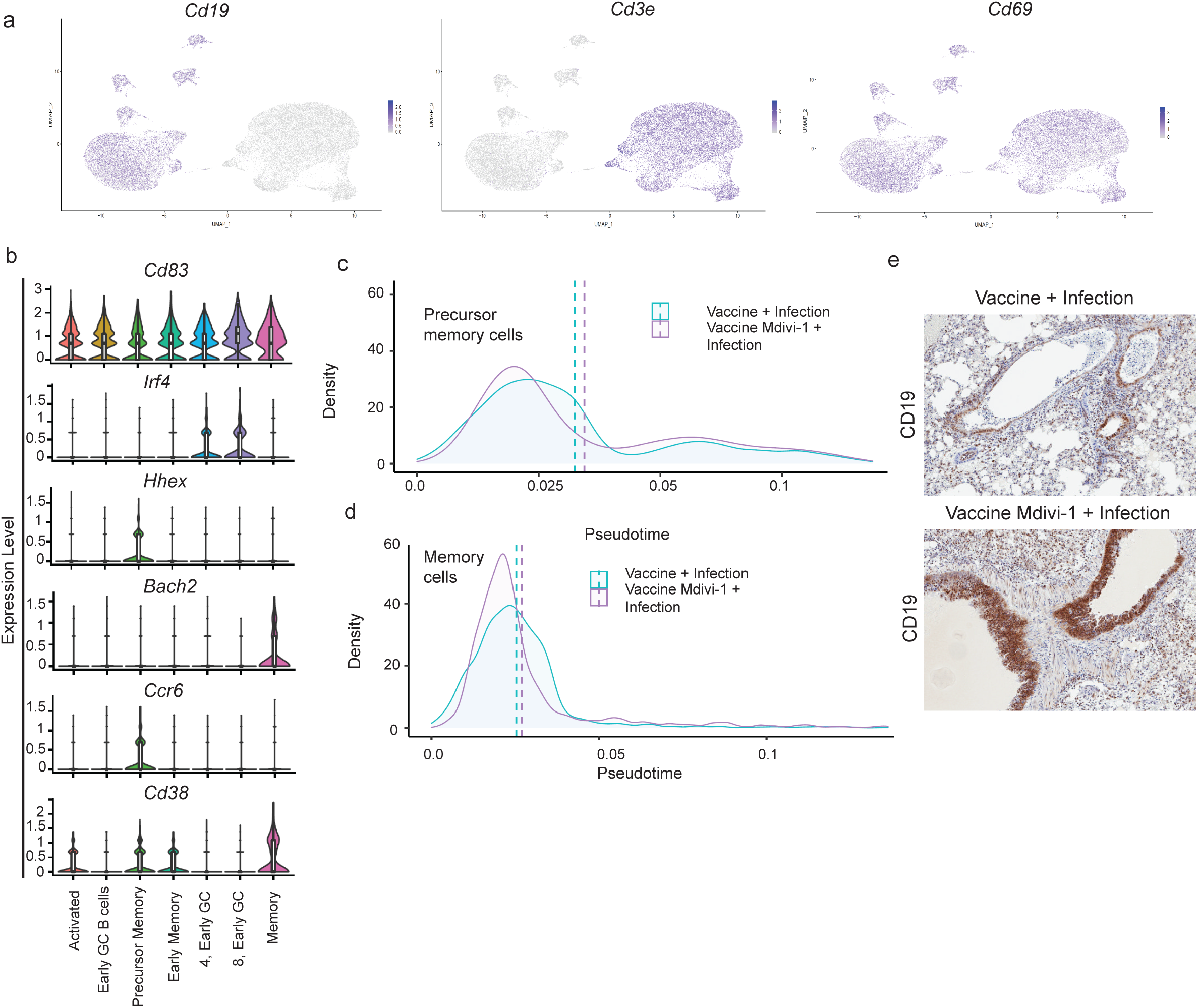
Mitochondria fission inhibitor supports memory B cell differentiation during immune response and post infection *in vivo*. Markers for defining B cell clusters. a) UMAPs identifying different immune cells clusters. Cd19 to identify B cells, Cd3e for T cells and Cd69 for activated immune cells. b) Violin plots of cells expressing respective genes from different B cell clusters to identify B cells populations among the clusters. Histograms (x-axis as pseudo-time and y-axis as normalized cell counts) depicting activated cells differentiation to c) lineage 2, Precursor Memory cells; d) lineage 3, Memory B cells in Vaccine + Infection and Vaccine (Mdivi-1) + Infection groups. e) Cd19 immunohistochemistry in mice lungs from Vaccine + Infection and Vaccine (Mdivi-1) + Infection groups.

**Supplementary Figure 4.**
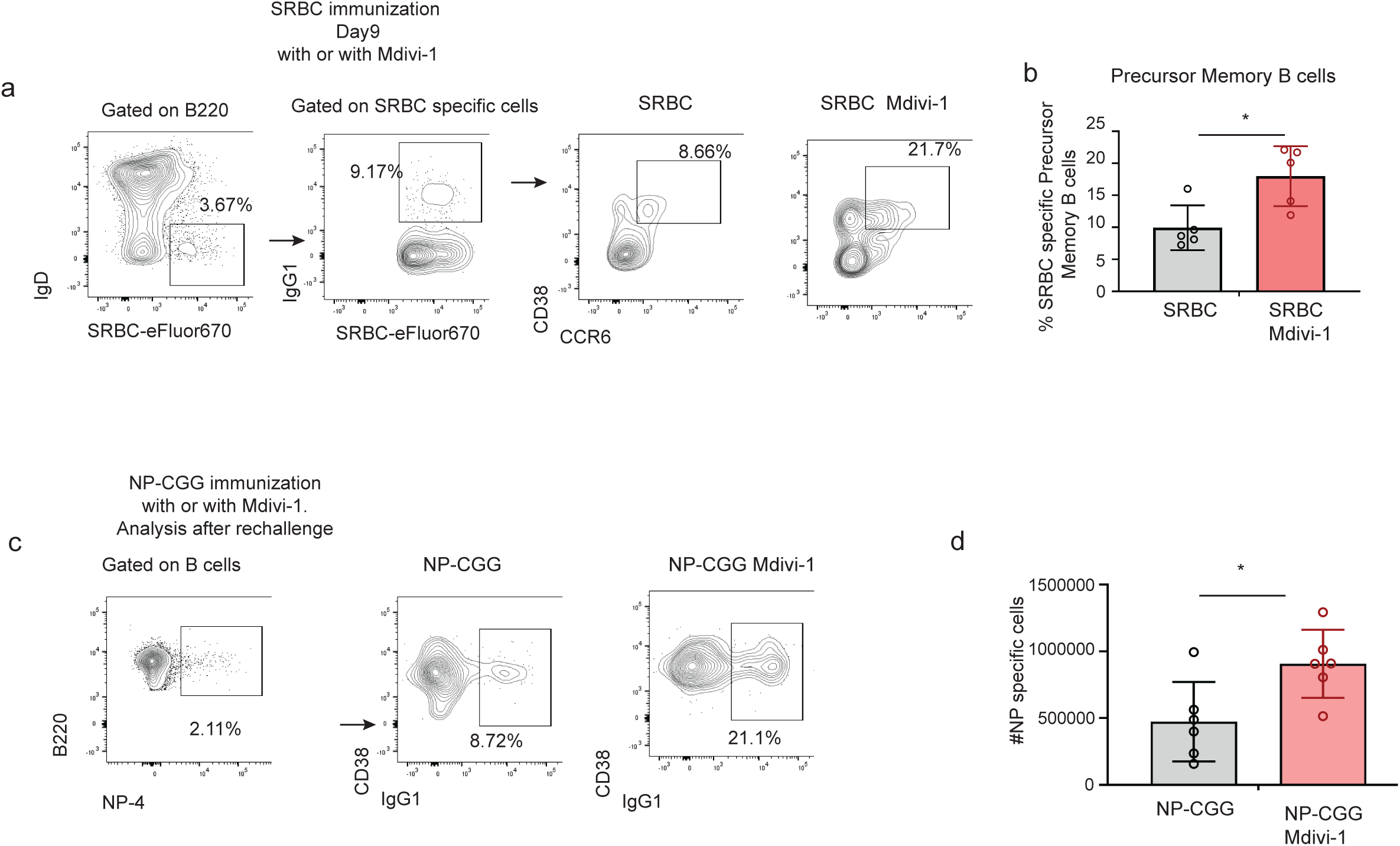
Mdivi-1 reinforces memory B cell differentiation during immune responses *in vivo*. Mice immunized with Mdivi-1 have enhanced precursor memory cells and replenishes their memory pool upon re-challenge. a) Gating strategy to identify SRBC-specific precursor memory B cells. Mice (n=5) each in group were immunized with SRBC on Day 0. Mice were given Vehicle or Mdivi-1 at 2.5mg/kg on Day 3 and Day 5. Spleens were harvested on Day 9 to study precursor memory B cells. b) Graph showing the SRBC-specific precursor memory cells in immunized mice with vehicle or Mdivi-1. c) Gating strategy for NP-4 specific memory B cells. d) Graph represents absolute number of NP-4 specific memory B cells in spleen of mice immunized with NP-CGG with or without Mdivi-1. Mice (n=6) in each group were immunized with NP-CGG with or without Mdivi-1 treatment. Mice were re-challenged with NP-CGG in PBS on Day 61 and were analyzed on Day 82. Statistical significance was calculated using t-test for b) and d); in prism p<0.01*.

**Supplementary Fig. 5.**
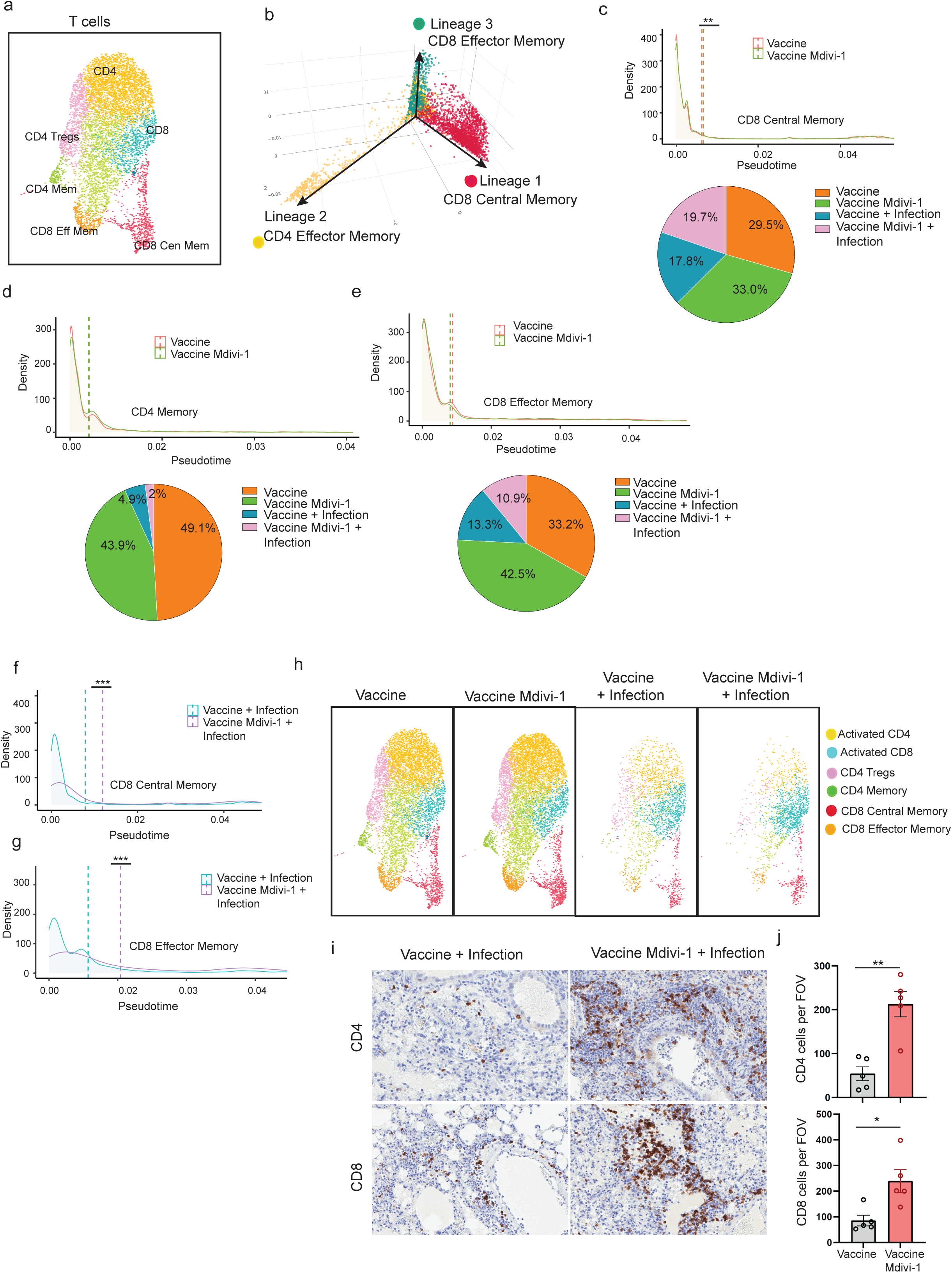
Mdivi-1 have subtle effect on memory T cells differentiation *in vivo*. a) UMAPs identifying T cells clusters in cells from all the samples. b) Displayed is the pseudotime lineage maps from B cells depicting activated cells differentiating into lineage 1, CD8 central memory, lineage 2, CD4 effector memory, lineage 3, CD8 effector memory. Histograms (x-axis as pseudotime and y-axis as normalized cell counts) depicting activated cells differentiation and below pi-charts showing T cells distribution in mice from Vaccine, Vaccine (Mdivi-1), Vaccine + Infection and Vaccine (Mdivi-1) + Infection group; c) lineage 1, CD8 central memory; d) lineage 2, CD4 effector memory and e) lineage 3, CD8 effector memory in Vaccine and Vaccine (Mdivi-1) treated mice. f) CD8 central memory, g) CD8 effector memory in Vaccine + Infection and Vaccine (Mdivi-1) + Infection group. h) UMAPs, identifying distinct T cell clusters showings T cells distribution in mice from Vaccine, Vaccine (Mdivi-1), Vaccine + Infection and Vaccine (Mdivi-1) + Infection group. i) Immunohistochemistry for CD4 and CD8 in mice lungs sections from Vaccine + Infection and Vaccine (Mdivi-1) + Infection group. j) Graph to quantify CD4 and CD8 positive cells in focus area. Statistical significance was calculated using Wilcoxon signed rank test for c-e, f-g, chi-square test for c-e, t-test for j in prism. p=0.04*, p=0.005**, p=0.0001***, p<0.0001****.

**Supplementary Fig. 6:**
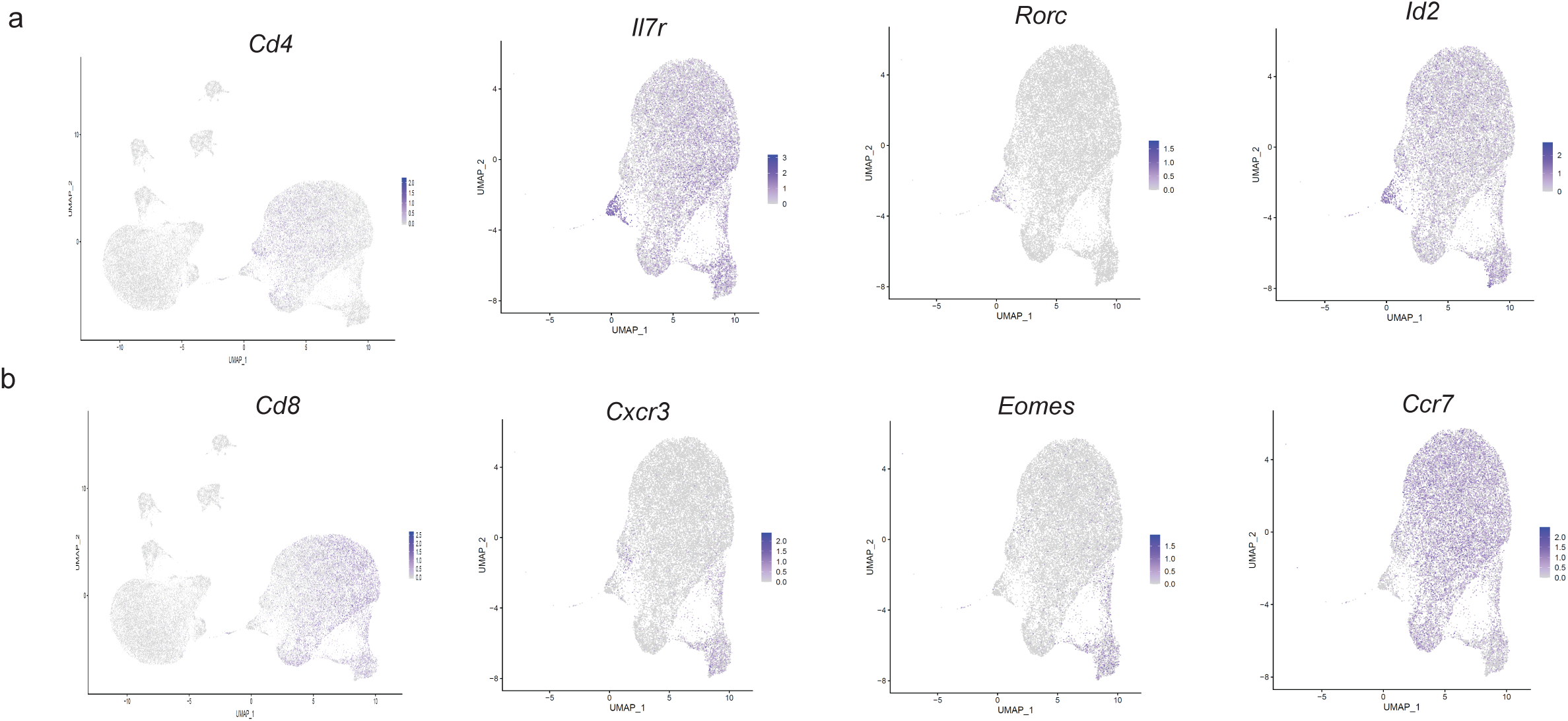
Identification strategy for T cell subsets clusters. UMAPs highlighting a) *Cd4*, *Il7r*, *Rorc* and *Id2*; b) *Cd8*, *Cxcr3*, *Eomes* and *Ccr7* expression in T cells from scRNA sequencing of mediastinal lymph nodes.

**Supplementary Fig. 7:**
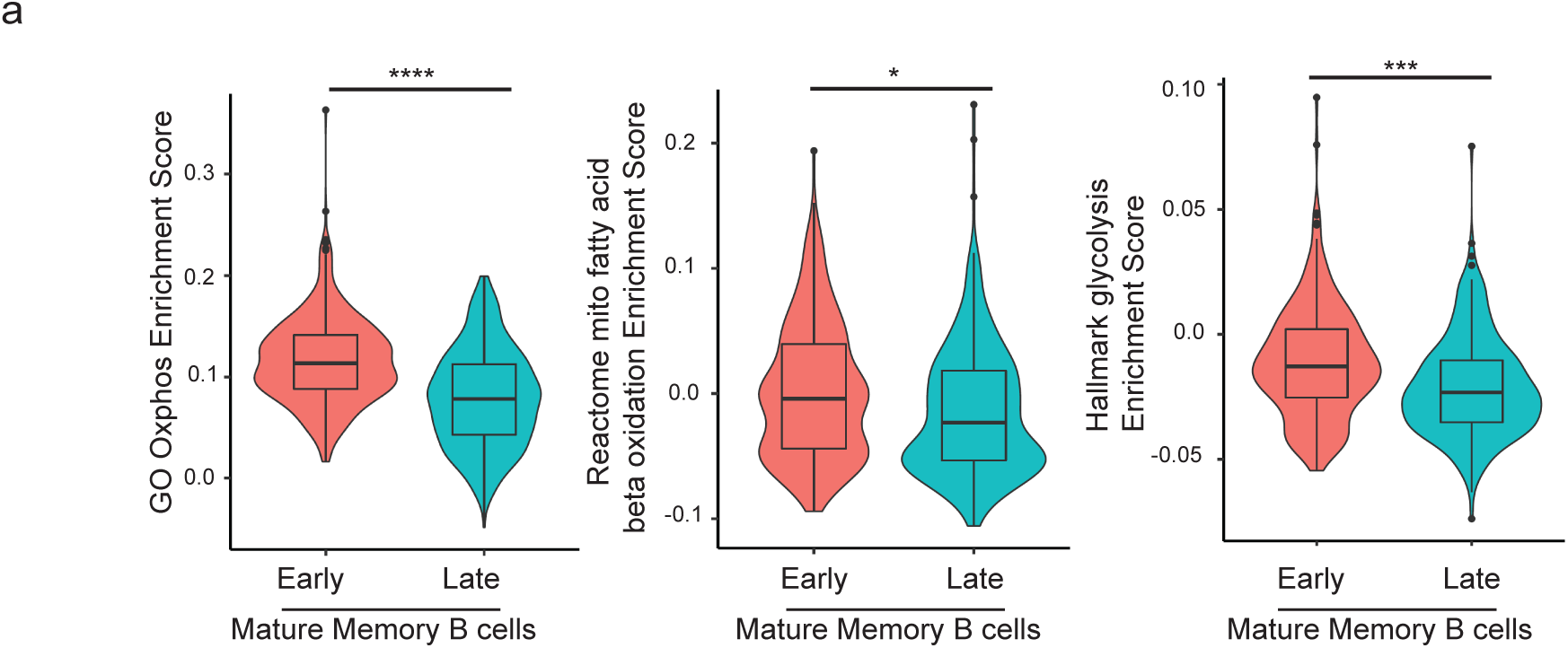
Mitochondrial pathways enrichment in memory B cells lineage. a) Normalized enrichment score of GO oxphos (left), Mitochondrial Fatty acid beta oxidation (center) and Hallmark glycolysis (right) pathways of the top and bottom 10% of grouped B cells by pseudotime and labelled the groups as “early” (start of the lineage 3) and “late” (end of the lineage 3). Statistical significance was calculated using linear model for K, p=0.04*, p=0.005**, p=0.0001***, p<0.0001****.

**Supplementary Fig. 8:**
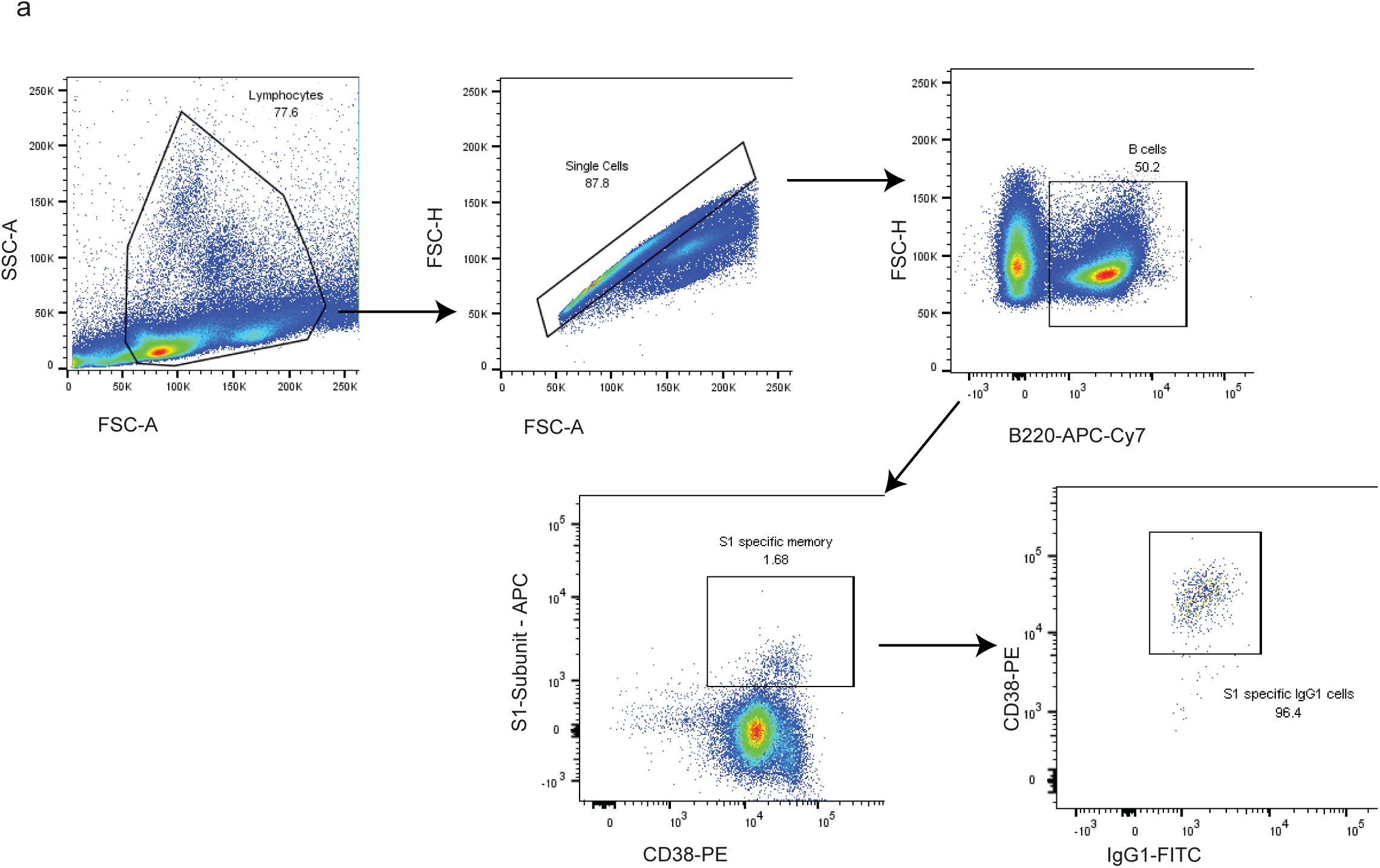
Gating strategy for identifying S1 subunit specific B cells. a) Gating scheme to identifying memory B cell and S1-subunit specific B cells in immunized mice.

## Methods

### Mice

All mice used in this study were on the C57BL/6J genetic background. Mice were either purchased from Jackson Laboratory or in-house vivarium at SBP. All mice used in the studies were 6-16 weeks of age and were housed in specific-pathogen free animal facility of Sanford Burnham Prebys Medical Discovery Institute. All the procedures were performed under to approved protocols by Institutional Animal Care and Use Committee (IACUC).

### Immunizations

For 4-hydroxy-3-nitrophenylacetyl-conjugated chicken gamma globulin, (NP-CGG, BIOSERACH, catalog # N-5055C-5) immunization, the hapten conjugated protein was diluted in 1mg/ml concentration (50 µl PBS NP-CGG per mouse) was mixed with 50 µl of Imject Alum Adjuvant (ThermoFischer Scientific, catalog number 77161). 100µl of solution was intraperitoneally injected to mice. For sheep red blood cells (SRBCs) immunization, citrated SRBCs (catalogue# 31102) were purchased from Colorado Serum Co, Denver, CO. SRBCs were two times washed with PBS. Washed SRBCs were resuspended 1:10 in PBS, mice were immunized with 100 µl on Day 0 and Day 5. For S1-Subunit, Recombinant SARS-CoV-2, S1 Subunit Protein (RBD) was purchased from RayBiotech, Code number-230-30162-100. 5µg of recombinant protein in 50µl was mixed with 50µl of Imject Alum Adjuvant overnight on rotor. 100µl of solution was intraperitoneally injected to mice.

### Mdivi-1 injections

Mdivi-1 was purchased from Cayman Chemicals (item no. 15559). The stock solution of Mdivi-1 was diluted in PBS to obtain 2.5mg in 100 µl. The drug solution was briefly sonicated for 10 seconds and passed through 0.22-micron filter. 100 µl of filtered drug solution was injected per mouse. Vehicle only was used as controls.

### Mitochondrial stress test

Splenic B cells were isolated by CD43 negative selection. Cells were incubated in media for 2 hours. Cells were cultured in complete Roswell Park Memorial Institute 1640 (Cellgro; Corning) supplemented with 10% FBS (Sigma-Aldrich), 1× penicillin/streptomycin (Cellgro; Corning), 2 mM GlutaGro (Cellgro; Corning), 1× MEM nonessential amino acids (Cellgro; Corning), 1 mM sodium pyruvate (Thermo Fisher Scientific), and 50 mM β-mercaptoethanol (Thermo Fisher Scientific). Cells were stimulated with anti-mouse Cd40 (clone: 1C10; Thermo Fisher Scientific) (1:200) and anti-mouse IgM F(ab′)_2_ (Jackson Immuno-Research) (5µg/ml) for 24 hours. After 24 hours molecules were added, M1 fusion promoter (20µM) (Sigma), Rapamycin (100nM) (SelleckChem), Mdivi-1 (10µM) (Cayman), Dynasore (10µM) (Cayman), Dyngo-4a (10µM) (Cayman), Hemin (60µM) (Sigma) and Bafilomycin (10nM) (Invivogen). At 48 hours, mitochondria stress test was performed using Aligent Seahorse XFe24 Analyzer to determine the oxygen consumption rate, cells were plated in triplicates. DMSO (Sigma) treated stimulated cells were used as control.

### Isolation of GC and Memory B cells

GC and memory B cells were negatively selected by MACs separation. For GC B cells, splenic B cells were stained for 30 mins with biotinylated antibodies for CD43, CD38, IgD and memory B cells CD43, IgD, Gl7. Cells were washed and incubated in biotin-binding streptavidin beads for 20 mins. For enriched GC and memory B cells, cells were passed through LD columns from Miltenyi.

### Confocal imaging

GC B cells were incubated on glass slides pre-coated with poly-l-lysine for 15 mins. Cells were fixed with 4% PFA for 10 mins at RT followed by two washes with PBS for 5 mins. Cell were next permeabilized with 0.5% Triton X-100 for 10 mins. Permeabilized cells were washed again two times with PBS for 5 min and are then blocked with 500 μl 1% BSA in PBS for 30 min. The blocking solution is replaced with 250 μl of the Tomm20 (Cell Signaling) 1:500 diluted in 1% BSA and left to incubate for overnight at 4°C. After 3 washes cells were incubated in secondary antibody anti-rabbit (PE) 1:2000 in 1% BSA for 1 hours at room temperature. Cells were counter stained with DAPI for 10 mins, washed 3 times with PBS. A drop a anti-fade fluorescence mounting medium was added and cover slip was placed. Slides were imaged on Nikon N-SIM E Super-Resolution/A1 ER Confocal Microscope system at SBP imaging core facility.

### Cas9 RNP targeting

Alt-R crRNA and Alt-R tracrRNA (from IDT) were reconstituted to concentration of 100 uM in Nuclease-Free Duplex buffer (IDT). RNA duplexes were prepared by mixing oligonucleotides (Alt-R crRNA and Alt-R tracrRNA) at equimolar concentrations in a sterile PCR tube (e.g. 4 uL Alt-R crRNA and 4 uL Alt-R-tracrRNA). Mixed oligonucleotides were annealed by heating at 95C for 5 minutes in a PCR machine, followed by incubation at room temperature for at least 1 hour. 5.9 uL of crRNA-tracrRNA duplexes (180 pmol) were then mixed with 2.1 uL IDT Cas9 v3 (80 pmol) (together Cas9 RNP) by gentle pipetting and incubated at room temperature for at least 10 min. 35 uL of cell media was aliquoted in a 24 well plate. 2 million B cells were then washed with PBS and resuspended in 15 uL resuspension buffer (4D-Nuclcofector X Kit S, #V4XP-4032; Lonza). Cells in resuspension buffer (12.5ul) were then mixed with Cas9-RNPs (7.5 ul) and transferred to the nucleofection cuvette strips and electroporated using the CM-137 program with 4D nucleofector. After electroporation, cells were transferred into 35 uL media in 24 well plate, and incubated for 20 min at 37C before transferring cells to activation media with 10ug/ml anti-mouse IgM and 200ng/ml anti-mouse CD40. *Drp*1 (Dnm1l) Antisense: 5’ AATCGTGTTACAATACTCTG 3’ *Cd4*: 5’ TATCACGGCCTATAAGAGTG 3’

### Influenza infection

All work with H1N1 virus was conducted in BSL-2 and Animal BSL-2 facilities under the protocols approved by SBP institutional board committee (IBC) and IACUC. The H1N1 (A/California/07/2009) strain of influenza virus was purchased from ATCC and produced in the MDCK cells. Mice were inoculated via intranasal route during light isoflurane (Aesica Queenborough Ltd., Kent, UK) anesthesia with 5×10^4^ plaque-forming units (PFUs) of virus in 50 μl of PBS for lethal infection dose. For vaccine, 2.5×10^2^ PFUs of H1N1 virus was heat inactivated at 90C for 10 mins and were administered to mice intranasally.

### Flow cytometry

Cells were stained in FACS buffer (0.5% bovine serum albumin, 2mM EDTA, and 0.05% sodium azide in PBS) with indicated antibodies for 30 mins on ice. Cells were washed and then fixed with 1% paraformaldehyde (diluted from 4% with PBS; Affymetrix) before FACS analysis using FACS Canto, X-20 and FACS LSR II (BD Biosciences). Antibodies and dye were from Thermo Fischer, ebiosciences, BioLegend, Southern Biotech and BD PharMingen. Data were analyzed with FlowJo (FlowJo LLC, Ashland, OR). Surface staining of cell suspensions was performed by treating the cells with anti-CD16/32 (clone 24G2; BD Biosciences), followed by staining with these antibodies: B220 (RA3-6B2), IgM (II/41), Fas (Jo2), GL7, CXCR4 (L276F12), CCR6 (29-2L17), IgD (11-26c), IgG1 (A85-1), anti-histidine (J095G46), MitoTracker Green FM (cat# M7514), MitoTracker Red CMXRos (cat# M7512). For recombinant SARs-CoV2 S-1 specific B cells cells were incubated with S1 subunit followed by secondary anti-histidine antibody.

## Serum antibody assays

NP-CGG immunization: Plates were coated overnight at 4°C with 50µl/well NP2 BSA at 10µg/ml diluted in PBS. Plates were next blocked by adding 200ul of blocking solution (1% BSA, 0.05% sodium azide in PBS) on top for 2 hours at 37°C. Plates were washed 2 times with DI water and 1 time with 200 μl of wash buffer (TBS containing 0.2% Tween 20). Next, 50ul diluted serum samples per well were added and incubated for 2 hours at 37°C. Plates were washed and incubated with 100 µl phosphatase substrate solution (Sigma) for 10–15 min, and absorbance was measured at 405 nm.

### S1-subunit ELISA

Plates were coated overnight at 4°C with 50µl/well of 4 μg/ml Recombinant SARS-CoV-2, S1 Subunit Protein (RBD) in PBS. Plates were washed 3 times with 200μl of wash buffer (TBS containing 0.2% Tween 20) and blocked with 100 μl of blocking buffer (3% milk in TBS containing 0.05% Tween 20) for 1 hour at 37°C. After removing blocking, 50μl of serum diluted in PBS was added and incubated for 2 hours at 37°C. The plates were washed in the wash buffer, 50 μl of alkaline phosphatase-conjugated secondary goat anti-mouse secondary antibody at 1:2500 dilution was added for 1 hour at 37°C. Plates were washed and incubated with 100 µl phosphatase substrate solution (Sigma) for 10–15 min, and absorbance was measured at 405 nm.

### Neutralization assay

ACE-2 receptor expressing cells Vero cells and luciferase expressing SARS-Cov2 pseudovirus particles were generously provided to us by Dr. Sumit Chanda’s group at SBP. To form antibody-virus complexes serum samples were incubated at 37°C with 5% CO_2_ for 1 hour. Neutralizing antibody serum samples were tested at a starting dilution of 1:10, and were serially diluted to 1:100 and 1:1000. Following incubation, growth media was removed and virus-antibody dilution complexes were added to the cells in duplicate. Virus-only controls and cell-only controls were included in each neutralization assay plate. After 48 hours cells were lysed and luciferase activity was measured by Nano-Glo Luciferase Assay System (Promega) according to the manufacturer’s specifications.

### scRNAseq data: quality control, normalization, and visualization

Mediastinal lymph nodes (mLNs) from mice n=5 in each group were pooled for each sequencing run. MLNs were homogenized in 70-µm cell strainer to obtained single cell suspension. Cells were shipped to Genewiz for sequencing. Quality control was performed on each individual single-cell sample. Low-quality cells were excluded from downstream analysis on the basis of having outlier mitochondrial content (indicative of cellular stress or damage), features counts or gene counts, using the R package Seurat (v3.1.5)(*1*, *2*). Outliers were determined as cells having more than the sample median + 5 median absolute deviations (mad) for each feature (mitochondrial content, feature counts, and gene counts). The highest upper threshold for mitochondrial gene content was 7%. The stage in cell cycle (e.g. S phase, G2M phase) was determined using Seurat’s built in function CellCycleScoring, with the cell cycle associated genes converted to their mouse homologs. Finally, since we detected a significant number of CD4/CD8 positive cells, we applied the R package DoubletFinder (v2.0.2)(*3*) with default parameters and found a preferential exclusion of CD4/CD8 positive cells indicating these cells were doublets. Cells passing quality control were merged and normalized together using SCTransform, with parameters; variable.features.n = 3,000 and regressing unwanted variance associated with mitochondrial content and cell cycle score (v0.2.0)(*4*). Cells were clustered using an SNN modularity optimization method on the first 10 principal components (using FindNeighbors and FindClusters functions from Seurat). The data was visualized using a Uniform Manifold Approximation and Projection (UMAP) projection. Cell types were identified first on the basis of expression of known immune cell type marker genes and also by comparison to built-in immune reference datasets (Encode and HPCA) using SingleR (v1.0.1)(*5*). We used Seurat’s FindMarkers function to identify cluster markers.

### Trajectory Analysis

To investigate lineage changes associated with Mdivi-1, we employed the trajectory inference method Slingshot (v1.3.1)(*6*). Slingshot analysis was separately performed on T cells and B cells, which were first subset and reclustered from the total cell dataset. To identify lineages, first we created a three-dimensional (3D) diffusion map embedding using the DiffusionMap function from the R package destiny (v3.2.0)(*7*). Lowly expressed genes we excluded when creating this embedding, only genes with at least 3 counts in 2 or more cells were retained. Slingshot was then applied unbiasedly (i.e. not pre-defining a starting cluster, or ‘root’) to the 3D diffusion map embedding. Three-dimensional visualizations were created using R package plotly (v4.9.2.2)(*8*). Slingshot was also applied in an unbiased manner to the UMAP embeddings to confirm the lineages identified in the diffusion maps. To determine genes significantly correlated with the slingshot-derived pseudotime values of each lineage, we used tradeSeq (v1.2.01)(*9*). Finally, to determine whether any of the lineages were associated with changes in pathway enrichment, we determined the average expression of genes from our pathways of interest using Seurat’s built-in AddModuleScore function. To identify pathways which were significantly correlated with increasing or decreasing pseudotime, a linear model was fit.

### Statistics

Data were analyzed in Prism 9 (GraphPad). Statistical tests were performed as indicated in the figure legends, and *n* values are also provided. All error bars represent mean ± s.d. Mice and tissues were randomly assigned to treatment groups where applicable. No data were excluded. Data were presumed to be normally distributed. Statistical significance was defined as *P* < 0.05, for reasons of space and visibility, the individual *P* values have not been integrated into the figures.

